# Evaluation of Anthropogenic influence on surface water quality and its ecological risk in the Ahafo Ano Southwest District

**DOI:** 10.64898/2026.07.30.741919

**Authors:** David Azanu, Afia Sarpong Anane Gyebi, Simeon Makafui Nutsuklo, Sampson Yaw Marfo, Isaac Ayew Aidoo, Francis Nyarkoh, Evans Barimah, Seifu A Tilahun, Junias Adusei-Gyamfi

## Abstract

Surface water quality is a significant public health concern because contaminated rivers and lakes can spread waterborne diseases and expose communities to harmful pollutants, particularly in areas where water treatment and regulation are limited. This study assesses the anthropogenic impact on surface water quality in the Ahafo Ano Southwest District, Ghana. Water samples were collected from three locations within the Mankran watershed during both wet and dry seasons and analyzed for key parameters. Pollutant concentrations exhibited marked spatial and seasonal variability, with nitrate ranging from 1.09 to 13.77 mg/L, phosphate from 1.58 to 10.55 mg/L, and total suspended solids from 140.0 to 1680.0 mg/L. Heavy metals frequently exceeded acceptable limits, with zinc ranging from below detection to 0.11 mg/L, copper from below detection to 0.84 mg/L, and lead from below detection to 0.46 mg/L. Seasonal Water Quality Index (WQI) analysis indicated degraded water quality during the wet season in Baniekrom and Kunsu, due to increased runoff carrying agricultural residues, sediments, and mining contaminants. In contrast, Mmobrem showed improved wet-season quality, attributed to rainfall dilution. The Piper diagram further revealed that the water samples predominantly fall within the Ca–Cl hydrochemical facies, reflecting strong geogenic control through basaltic rock weathering, coupled with chloride enrichment likely intensified by agricultural and domestic inputs. Principal component analysis indicated that multiple anthropogenic and natural factors drive water quality variations across the catchment. Ecological risk assessment identified potential risks to algae, resulting from elevated levels of nutrients and metals. Overall, the combined WQI and hydrochemical evidence indicate an intensified nutrient influx during the wet season and increased suspended particulate matter during the dry season, primarily driven by agriculture, deforestation, and mining. The findings underscore the urgent need for sustainable water management strategies to mitigate the impacts of human activities on surface water quality in the region.

## 1. INTRODUCTION

Surface waters account for only 2.5% of the world’s freshwater supply, which is essential for human survival and the health of ecosystems [1]. Surface water sources, including rivers, lakes, and streams, are crucial habitats for a wide range of aquatic plants and animals. They form the basis of complex food webs and contribute significantly to ecological health [2]. However, pollution and unsustainable activities put pressure on these water sources, resulting in water scarcity, ecological habitat destruction and the spread of waterborne infections in humans and animals [3].

Water pollution is pervasive in developing countries, with numerous studies focusing on the quality of surface water. For instance, Atiku et al.[4], conducted an in-depth investigation using biomonitoring techniques to assess river health, where the proliferation of bio-plants in water is indicative of pollution, and the seasonal variations in physicochemical parameters and chlorophyll contents in water bodies. Moreover, Toure et al. [5] examined the concentrations of heavy metals and their impact on surface water quality. Obasi et al. [6] explored the role of certain bio-flora species in aquatic environments, suggesting that these plants can act as pollutants. Similarly, Elshobary et al. [7] identified the presence of algae as a signal of water pollution, noting that specific groups of algae thrive in environments contaminated with organic matter and sewage. Researchers such as Atiku et al. [4]and Obasi et al. [6], have used algal growth as a bioindicator for assessing water quality while Elshobary et al. [7] employed diatoms to determine water quality standards. The physicochemical and biological characteristics of water are affected by industrial discharges, thereby impacting water quality. As carbon dioxide increases, the concentration of dissolved oxygen (DO) and phosphate decreases, reducing waste content in water. Aniyikaiye et al. [8], evaluated physicochemical characteristics of water, such as pH, chlorine, TDS, conductivity, and others. The study compared treated and untreated water samples and resolved that the quality of water increases after treatment. Williams et al. [9] assessed water quality in several water samples using physicochemical analytic tools and proposed that water quality must be improved and treated well before consumption.

Anthropogenic activities often exacerbate water pollution in developing countries. Agriculture, a primary economic sector, substantially contributes to water pollution through the runoff of fertilizers and pesticides [10]. Water can be polluted by runoff from agricultural activities, industrial discharges, and untreated human and animal waste. Aside from pollution that is human-induced, contamination can also be caused by natural means [11]. The seepage of chemicals like arsenic or fluoride into drinking water caused contamination. In the African sub-region, many water bodies are contaminated. Human and animal excreta primarily cause this situation in Africa, primarily through human defecation that dissolves in water. Microbiological contaminations are caused by bacteria, viruses and protozoa, which are collectively referred to as pathogens [7].

Ghana with abundant surface water resources including three main Basins namely the Volta Basin, Southwestern Basin (Ankrobra, Pra, and Tano), and the Eastern-Coastal Basin (Densu) according to the Water Resources Commission (WRC) of Ghana [12]. The management and sustainable utilization of these resources are essential for ensuring the socio-economic development and well-being of the Ghanaian population [12]. However, the quality of surface water is increasingly threatened by anthropogenic activities, leading to significant challenges for water availability and accessibility [13].

The Ahafo Ano Southwest District is a perfect example of how human activities, water resources, and environmental risks are all connected. With roughly 65,770 residents as of 2021, the district makes a living mainly from farming, especially growing cocoa and coffee as well as small-scale mining. Despite the abundance of surface water resources, including the Tano River and numerous streams and ponds, the district faces growing concerns regarding its sustainability [14]. The land of Ahafo Ano South in Ghana is rich in gold, attracting miners seeking their fortune. However, this pursuit has come at a cost, impacting the very lifeblood of the region. One of the main concerns is the widespread use of mercury in small-scale or artisanal gold mining. Mercury is highly toxic, and when it enters the water through improper disposal or accidental spills, it contaminates the entire ecosystem [15]. This contamination not only harms fish and other aquatic life, but also poses a serious health risk to communities who rely on the rivers for drinking water, cooking, and even bathing.

This research aligns with Sustainable Development Goals (SDGs) 6 and 13, which aim to ensure access to clean water and sanitation for all, and to combat climate change, respectively. It also supports national policies, such as the Ghana Water Sector Strategic Development Plan and the National Water Policy, which emphasize universal access to safe water and the sustainable management of water resources. This study aims to promote targeted interventions, including sustainable farming practices and improved sanitation, to protect the district’s water resources and environment by identifying the primary factors contributing to water pollution. This study aims to assess the impact of human activities on the surface water resources of the Ahafo Ano South West District, Ghana. This will be achieved by assessing both the water quality and quantity within the district, identifying the human activities that most significantly contribute to water pollution, and ultimately determining the environmental risks associated with these anthropogenic influences.

## 2. MATERIALS AND METHODS

### 2.1 Study area

The target landscape for this study was the Mankran watershed in the Upper Offin Sub-basin of the Pra Basin (Fig. 1). The Upper Offin watershed spans an area of 3070 km², directly draining into the Offin River, whereas the Mankran watershed encompasses 122 km². The selected study sites for this monitoring initiative were identified to reflect various drivers of change within the micro-watershed. Kunsu exemplified the effects of illegal mining activities; Barniekrom highlighted the drivers linked to agricultural expansion; and Mmrobem illustrated the impacts of agricultural growth, extensive logging, and deforestation. These competing land uses exert unique influences on the water system, which need to be evaluated to ensure sustainable landscape management in the target district for both socio-economic and ecological advantages.

**Fig. 1.**
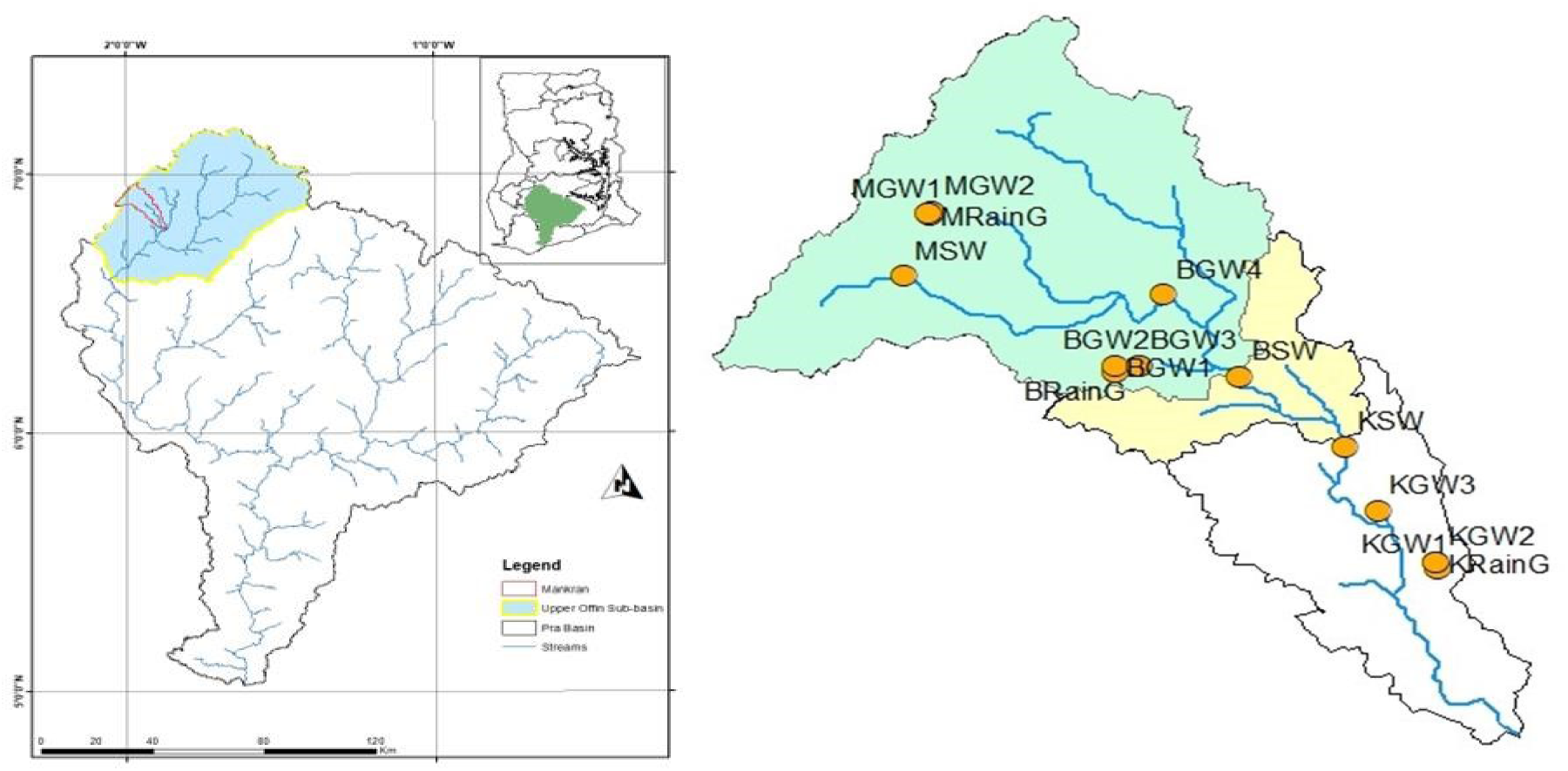
A map showing the study area and sampling points in the Mankranso watershed

The micro landscape of Mmrobem was primarily characterized by the presence of two significant drivers of change: agricultural expansion and deforestation. These two forces had far-reaching consequences on both the environment and the local communities. Agricultural expansion often involves the conversion of natural forest landscapes into farmland, which can lead to deforestation, soil erosion, and changes in land use. The monitoring program in the community covered one surface water location for water quality (MSW in Fig. 1). The primary driver of variations in hydrological quality and quantity was agricultural expansion. The monitoring program covered one surface water location for streamflow and water quality monitoring (BSW in Fig. 1). Kunsu sampling location represented the impact of illegal mining on hydrological variables. The monitoring program covered one surface water location for streamflow and water quality monitoring (KSW in Fig. 1).

### 2.2 Sampling

Water samples were collected using 500 mL polypropylene bottles from June 2023 to February 2024. At each sampling location, samples were taken in triplicate for physiochemical, heavy metals, and microbial analysis. Samples for heavy metal analysis were acidified with 5 mL of concentrated nitric acid (HNO_3_) and transferred to the laboratory for analysis in coolers at a temperature of 4 °C. Quality assurance and quality control (QA/QC) procedures were implemented throughout sampling and analysis to ensure data reliability. Duplicate samples were collected to assess analytical precision, while field blanks were used to check for potential contamination during sampling and transport. All instruments used for physicochemical and heavy metal analyses were calibrated using appropriate standard solutions prior to measurement, and calibration was verified periodically during analysis. Analytical procedures followed standard methods, and appropriate quality control checks were implemented to ensure data accuracy. All analyses were conducted following standard methods, and results were accepted only when quality control criteria were met.

### 2.3 Physicochemical analysis

The collected water samples were examined in accordance with the standards of the Environmental Protection Agency of Ghana and the United States Environmental Protection Agency (US-EPA).

Nitrate and phosphate were analyzed using the Agilent Cary 60 UV-Vis spectrophotometer (Agilent Technologies, Santa Clara, USA), The TB1 Velp Scientifica turbidimeter (VELP Scientifica Srl, Usmate Velate, Italy) was used to analyze the turbidity of samples. TDS, EC and Salinity were analyzed using an Ohaus starter 3100C bench meter (Ohaus Corporation, Parsippany, USA). pH was analysed using an Ohaus starter 3100 bench meter (Ohaus Corporation, Parsippany, USA).Chloride was determined by a Titrimetric principle (Mohr’s Method), and Total Suspended Solid (TSS) was determined by the gravimetric method.

### 2.4 Metals analysis

A 100 mL water sample was digested with 5 mL of nitric acid. It was heated for 1 hr at 250 °C. Metals were analyzed using Agilent 4210 Microwave Plasma Atomic Emission Spectroscopy (MP-AES) (Agilent Technologies, Santa Clara, USA).

### 2.5 Ecological Risk Assessment

According to Razak et al. [16], a risk quotient (RQ), which is the proportion of an element’s maximum measured environmental concentration (MEC) to its anticipated PNEC, was used to assess the potential harm of each metal presented to surrounding species. RQ was calculated at three trophic levels of the ecosystem: seaweed (*Desmodesmus subspicatus*), water flea (*Daphnia magna*), and fish (*Oncorhynchus mykiss*) to account for multiple stages of the stream’s dietary system. PNEC levels used in this risk evaluation are based on the lowest metal PNEC values found in the research or estimated using the ECOSAR system [17]. By multiplying the toxic effects data by an evaluation factor (EF), PNEC was determined. The expected value (EF) is a discretionary ratio used to account for the inherent uncertainty in the data of the metals collected in the research setting. For this study, PNECs were derived using an EF of 100 [18]. A commonly used risk rating approach also utilizes an RQ of less than 0.1, indicating the lowest hazard to freshwater organisms. An RQ of 0.1 or less and an RR greater than or equal to 1 indicates intermediate hazards, while an RR greater than 1 indicates extreme risk.

### 2.6 Water Quality Index (WQI) Determination

The Water Quality Index (WQI) was calculated to provide a single numerical indicator representing the overall suitability of the sampled water for domestic use. The WQI integrates multiple physicochemical parameters into a single value by applying weighting factors and quality ratings to each parameter [19,20].

Parameters included in the WQI assessment were those commonly used for surface and groundwater monitoring and recommended by international drinking-water guidelines. These included: pH, EC, TDS, Temperature, Turbidity, Nitrate, Phosphate, Chloride, Calcium, Magnesium, Sodium, and Potassium. The selection followed WHO/FAO recommendations for drinking-water assessment . The quality rating (Qi) for each parameter was calculated to express how closely the measured concentration aligned with its respective water-quality standard (Si). Surface water standards were used as reference values.

### 2.7 Statistical analyses

The mean, median, minimum, and maximum descriptive statistics for the concentration of PTEs were computed using Microsoft Office Excel 2019 (Version 25, Microsoft, USA). Principal correlation analysis, which identifies multivariate relationships, was conducted using GraphPad Prism version 10.4 for Windows (GraphPad Software Inc., USA). Eigenvalues greater than 1 were chosen based on the Kaiser rule. The calculated cation and anion percentages were plotted on a Piper trilinear diagram using Golden Software Grapher (Windows Version 25.3.315) software. The cation field was used to classify water types based on Ca²⁺–Mg²⁺, (Na⁺ + K⁺) distribution, while the anion field classified SO₄²⁻– Cl⁻– HCO₃⁻ facies (Piper, 1944). The central diamond field was then used to interpret integrated hydrochemical facies and categorize the groundwater samples into the following types: Ca–HCO₃, Ca–Mg–HCO₃, Na–Cl, Mixed Ca–Mg–Cl, and Mixed Na– HCO₃. The interpretation of water types was based on the clustering patterns of the samples within the main hydrochemical fields of the Piper plot [21,22].

## 3. RESULTS

### 3.1 Physicochemical Parameters

Table 1 summarizes the physicochemical parameters of the samples, including minimum and maximum values, as well as the World Health Organization-recommended limits for surface water. Water quality analysis across multiple locations and timeframes reveals significant variability in several parameters, some exceeding WHO-recommended limits for surface water, which raises environmental and health concerns. Zinc concentrations ranged from below detection limits to a maximum of 0.107 mg/L, substantially below the WHO limit of 3 mg/L, indicating no significant zinc contamination. Cadmium levels also remained low, peaking at 0.0024 mg/L, within the WHO acceptable limit of 0.003 mg/L, suggesting a limited presence of cadmium, likely influenced by minor anthropogenic activities or natural sources. In contrast, iron concentrations reached a maximum of 11.951 mg/L, far exceeding the WHO limit of 0.3 mg/L. This indicates significant contamination, potentially resulting from the natural leaching of iron-rich geological formations, industrial discharge, or the corrosion of metallic infrastructure. Copper concentrations varied, with a peak of 0.84 mg/L, which remains below the WHO limit of 2 mg/L but indicates localized variability, possibly due to runoff from agricultural activities or copper-containing industrial effluents. Nickel concentrations ranged up to 0.11 mg/L, exceeding the WHO guideline of 0.07 mg/L, highlighting potential contamination from industrial processes or electroplating activities. Lead levels exhibited concerning variability, with a maximum of 0.46 mg/L, exceeding the WHO limit of 0.01 mg/L, indicating severe contamination likely due to vehicular emissions, battery waste, or lead-based paints in the vicinity. Manganese concentration peaked at 1.33 mg/L, surpassing the WHO limit of 0.1 mg/L, which suggests that natural sources, such as leaching from soil or rocks, or contamination from mining activities, may be contributing to the elevated levels. Chromium concentrations also exceeded the WHO limit of 0.05 mg/L, with a maximum of 0.2 mg/L, possibly stemming from industrial discharges, including tanning or metal plating operations. Sodium and potassium levels varied widely, with peaks of 20.02 mg/L and 21.44 mg/L, respectively, within the acceptable ranges for surface water. These fluctuations might result from natural mineral dissolution or agricultural runoff. Magnesium and calcium levels, with maximums of 14.053 mg/L and 19.71 mg/L, respectively, were well below thresholds, indicating minimal concerns. Arsenic concentrations remained low, peaking at 1.11 µg/L, well within the WHO limit of 10 µg/L, while mercury levels (maximum 0.61 µg/L) also stayed within the WHO limit of 1 µg/L. Turbidity, however, exhibited alarming variability, reaching a maximum of 972 NTU, far above the WHO limit of 5 NTU. High turbidity could be attributed to suspended solids, sediment runoff from agricultural lands, or discharge of untreated wastewater. Nitrate levels were within safe limits, with a peak of 13.77 mg/L (WHO limit: 50 mg/L). However, phosphate concentrations peaked at 10.55 mg/L, indicating potential nutrient pollution from agricultural fertilizers or detergents, which can lead to risks of eutrophication. Total dissolved solids (TDS) and electrical conductivity (EC) values were within acceptable limits, but total suspended solids (TSS) peaked at 1680 mg/L, indicating significant suspended particulate matter, possibly from soil erosion or industrial discharge. Chloride and pH levels remained within acceptable ranges, ensuring minimal concerns about water salinity or acidity.

**Table 1.** Results of the water quality analysis compared to WHO WHO-recommended limits for surface water.

| Parameter | WHO limit for surface water (WHO, 2021) | Ghana EPA (2021) | Minimum | Maximum |
| --- | --- | --- | --- | --- |
| Zn(mg/L) | 3.0 | 3.0 | 0.000 | 0.107 |
| Cd(mg/L) | 0.003 | 0.003 | 0.000 | 0.002 |
| Fe(mg/L) | 0.3 | 0.3 | 0.000 | 11.951 |
| Cu(mg/L) | 2 | 2.0 | 0.000 | 0.845 |
| Ni(mg/L) | 0.07 | 0.07 | 0.002 | 0.114 |
| Co(mg/L) | - | 0.05 | 0.000 | 0.038 |
| Pb(mg/L) | 0.01 | 0.01 | 0.000 | 0.461 |
| Mn(mg/L) | 0.1 | 0.4 | 0.000 | 1.337 |
| Cr(mg/L) | 0.05 | 0.05 | 0.000 | 0.192 |
| Na(mg/L) | 200 | 200 | 5.111 | 20.017 |
| K(mg/L) | - | 12 | 0.927 | 21.438 |
| Mg(mg/L) | 50 | 50 | 3.955 | 14.053 |
| Ca(mg/L) | 75 | 75 | 3.162 | 19.710 |
| As(µg/L) | 10 | 10 | 0.000 | 1.108 |
| Hg (µg/L) | 1 | 1 | 0.000 | 0.610 |
| Sal (mg/L) | - | 1000 | 0.060 | 0.140 |
| pH | 6.5-8.5 | 6.5-8.5 | 7.030 | 7.910 |
| TDS (mg/L) | 1000 | 1000 | 60.400 | 125.400 |
| EC (µS/cm) | 1200 | 1500 | 120.800 | 251.000 |
| TSS (mg/L) | - | 25 | 140.000 | 1680.000 |
| Turbidity | 5 | 5 | 1.700 | 972.000 |
| Nitrate (mg/L) | 50 | 50 | 1.094 | 13.770 |
| Phosphate (mg/L) | - | 2.0 | 1.583 | 10.555 |
| Chloride (mg/L) | 250 | 250 | 7.490 | 129.900 |

### 3.2 Pearson correlation

The analysis of metal concentrations in the samples (Fig. 2) revealed several notable correlations, both positive and negative, with varying degrees of strength. In particular, a strong positive correlation was observed across the samples, indicating that as the concentration of one metal increased, the concentrations of others tended to rise as well.

**Fig. 2.**
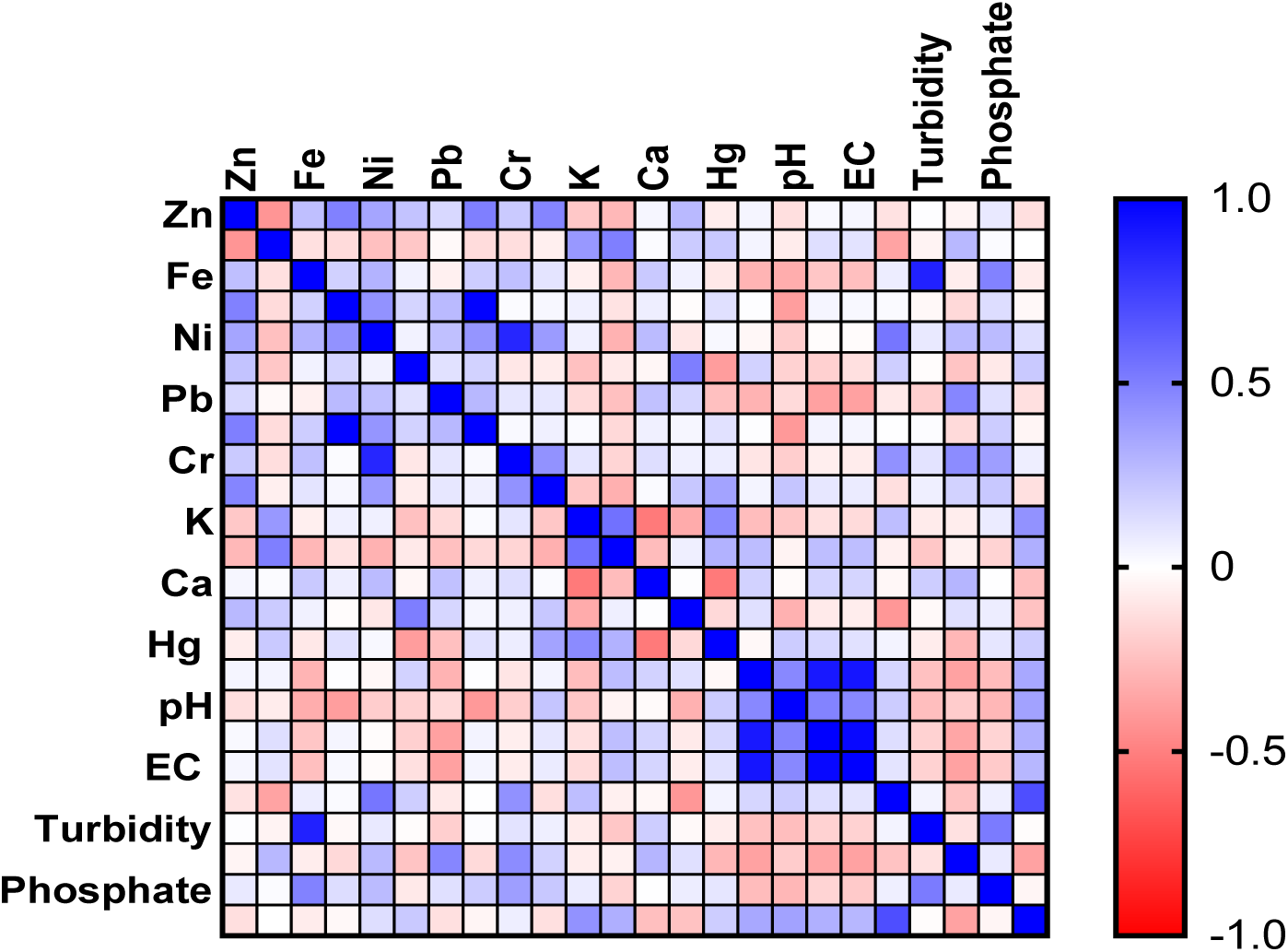
Pearson correlation heatmap

However, the relationship between zinc and cadmium deviated from this trend, displaying a strong negative correlation. This suggests that as zinc concentration increased, cadmium concentration decreased significantly, and vice versa. Additionally, a significant difference was observed between the concentrations of zinc and cadmium, underscoring the distinct relationship between these two metals. The correlation between nickel and copper (Fig. 2) was found to be weakly positive. Although the two metals tended to increase together, the relationship was not as pronounced as in other cases. Despite the weak correlation, a significant difference remained between the samples, indicating that the variations in their concentrations were statistically meaningful. Similarly, manganese and chromium showed a weak positive correlation, suggesting a slight tendency for their concentrations to increase together. As with the relationship between nickel and copper, the correlation was not particularly strong; yet, the differences in manganese and chromium concentrations across the samples were significant. In addition to these relationships, a weak positive correlation was identified between copper and total suspended solids (TSS). Although the correlation was weak, the statistical analysis indicated a significant difference in concentrations. Lead and pH also showed a strong negative correlation, although there was a significant difference between them.

### 3.3 Seasonal variation

Table 2 compares the average and standard deviation of analysed water quality parameters between the wet and dry seasons. Zinc concentration in the wet season (0.0348 mg/L) was slightly higher than that of the dry season (0.0273 mg/L). The dry season shows a much higher average TSS (946.89 mg/L) compared to the wet season (192.00 mg/L). The wet season recorded higher nitrate levels (7.14 mg/L) compared to the dry season (2.85 mg/L).

**Table 2:**
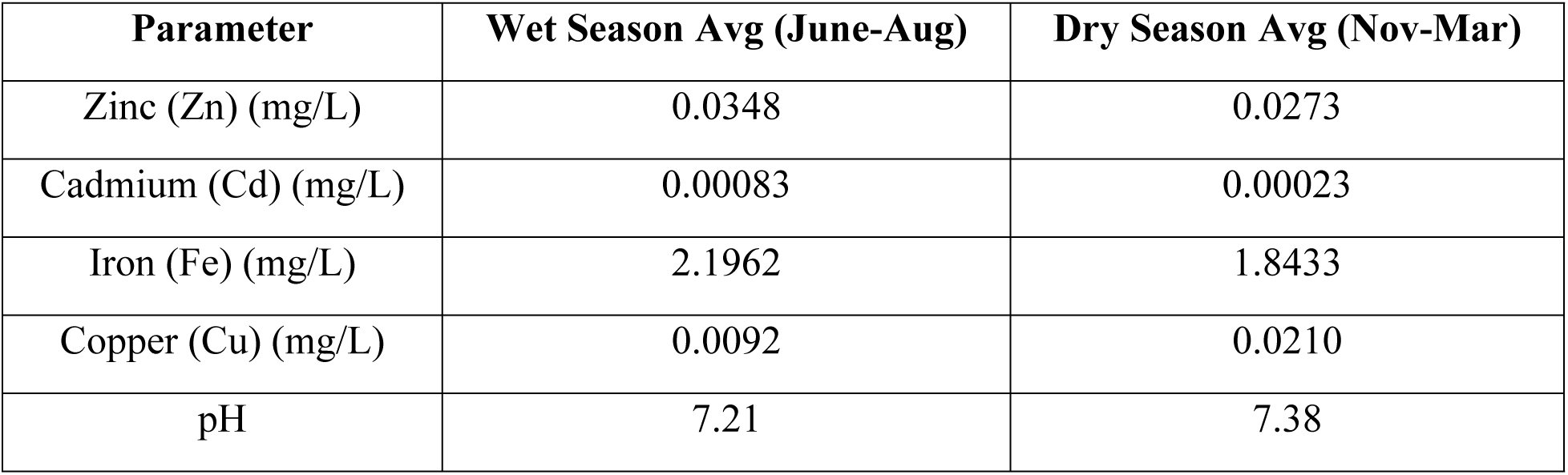
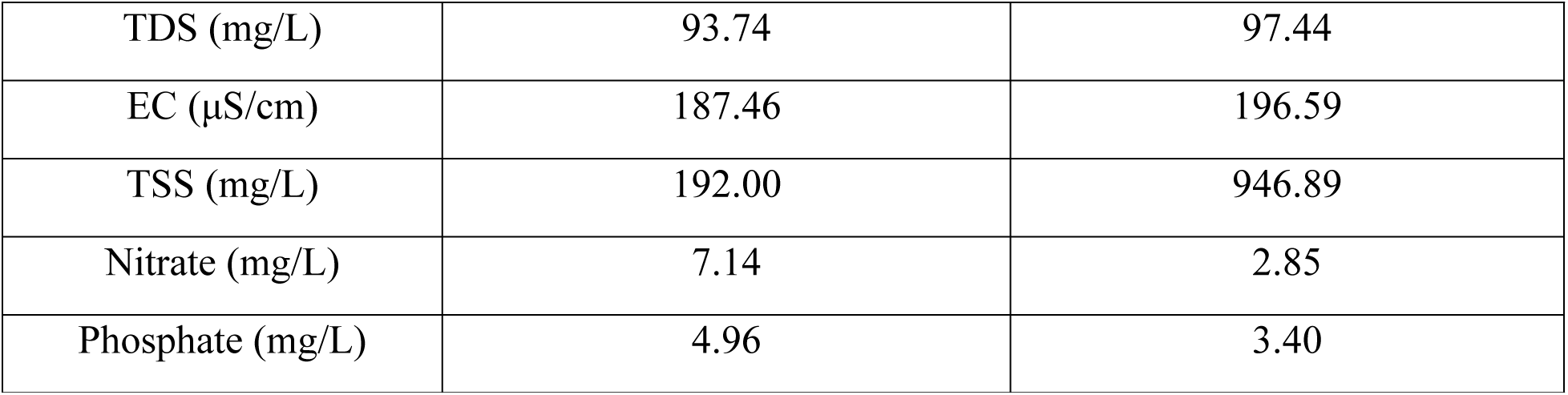
Average and standard deviation of water quality parameters between the wet and dry seasons.

| Parameter | Wet Season Avg (June-Aug) | Dry Season Avg (Nov-Mar) |
| --- | --- | --- |
| Zinc (Zn) (mg/L) | 0.0348 | 0.0273 |
| Cadmium (Cd) (mg/L) | 0.00083 | 0.00023 |
| Iron (Fe) (mg/L) | 2.1962 | 1.8433 |
| Copper (Cu) (mg/L) | 0.0092 | 0.0210 |
| pH | 7.21 | 7.38 |
| TDS (mg/L) | 93.74 | 97.44 |
| EC ($\mu\text{S}/\text{cm}$) | 187.46 | 196.59 |
| TSS (mg/L) | 192.00 | 946.89 |
| Nitrate (mg/L) | 7.14 | 2.85 |
| Phosphate (mg/L) | 4.96 | 3.40 |

### 3.4 Principal component analysis

The PCA biplot (Fig. 3) summarizes the multivariate relationships among the measured water quality parameters. All measured variables were looked at in terms of factor loadings to be able to interpret the underlying processes that affect water quality. Positive loadings that are strong (≥ 0.7) indicate the variables that make a strong contribution to one of the components, whereas moderate (0.500.7) and weak (<0.500.7) loadings represent the levels of the contribution. In this analysis, parameters of TDS, pH, Hg and chloride, showed high positive loadings on the first principal component (PC1), indicating a common source or process, probably related to mineral dissolution and anthropogenic inputs. On the other hand, nitrate and phosphate were negatively loaded, meaning that they were negatively correlated to the variables that predominated PC1, which could be due to the dilution of the nutrients or separate sources of pollution like agricultural runoff. The PC scores help identify groups or clusters of observations. The first principal component (PC1) is primarily associated with TDS, pH, Hg, and chloride, which load positively along this axis, indicating strong positive correlations among these variables. In contrast, nitrate and phosphate load negatively on PC1, suggesting an inverse relationship with the positively loaded parameters. The second principal component (PC2), captures additional variability. The distribution of sample scores (grey points) reflects similarities among sampling sites, where samples positioned closer together exhibit comparable water quality characteristics, while separation along PC1 and PC2 indicates differing physicochemical and nutrient profiles.

**Fig. 3.**
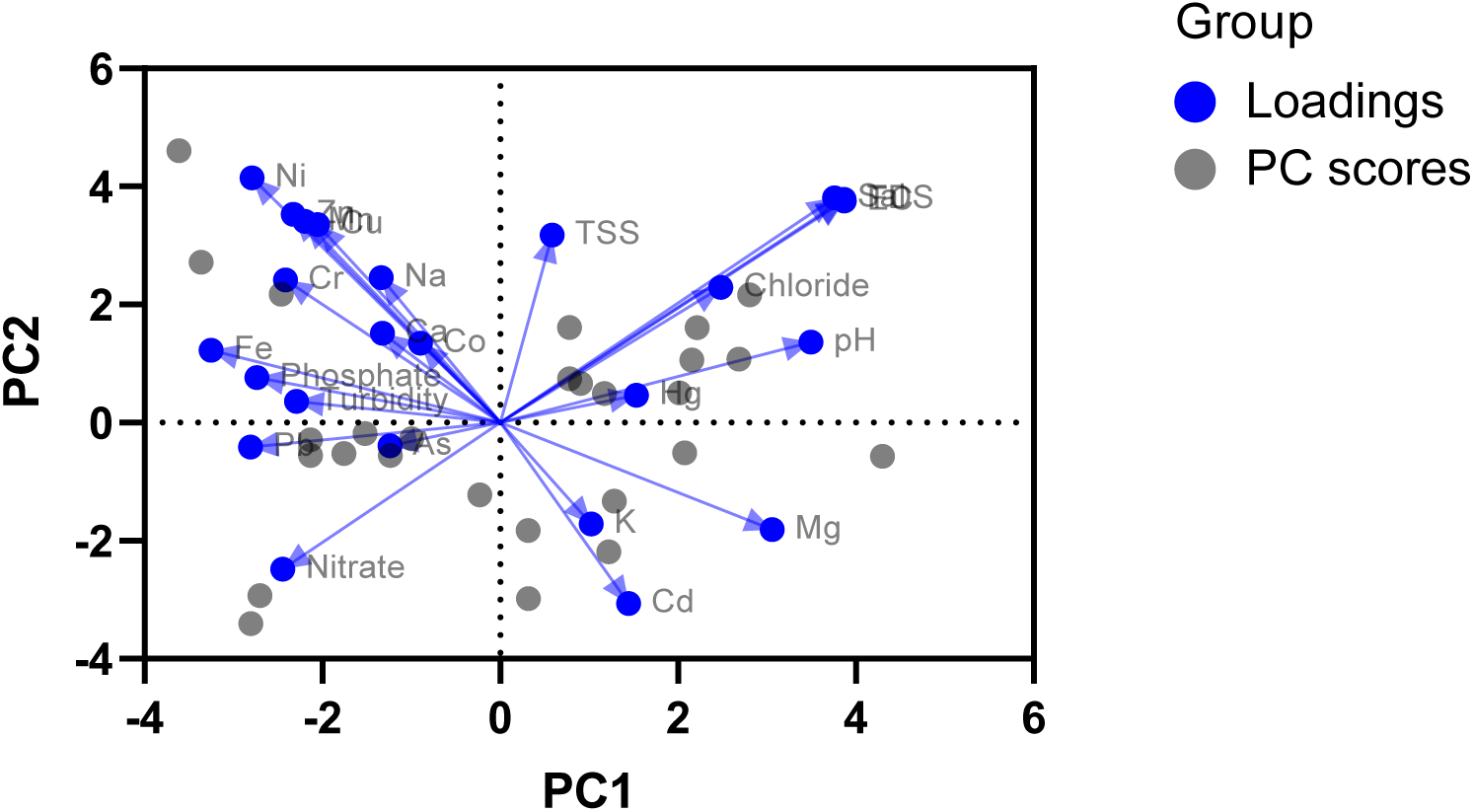
Biplot of principal component analysis

Most parameters showed seasonal variation: Zn, Cd, Fe, nitrate, and phosphate were higher in the wet season, likely due to runoff, while Cu, pH, TDS, EC, and especially TSS were higher in the dry season, reflecting reduced dilution and sediment concentration. pH remained neutral to slightly alkaline, indicating stable acid–base conditions.

### 3.5 Water Quality Index

The mean water quality indices (WQI) for Baniekrom were 97.46 in the wet season and 77.39 in the dry season, indicating a decline in water quality during the dry period. Monthly WQI values ranged from 298.17 to 3.31 (June–September) in the wet season and 199.08 to 1.50 (October– February) in the dry season (Fig. 4), reflecting substantial temporal variation. These values suggest that while water quality improves during the wet season due to rainfall dilution, some periods particularly in the dry season may render the water unfit for drinking without treatment, highlighting the need for continuous monitoring and appropriate water management practices.

**Fig. 4.**
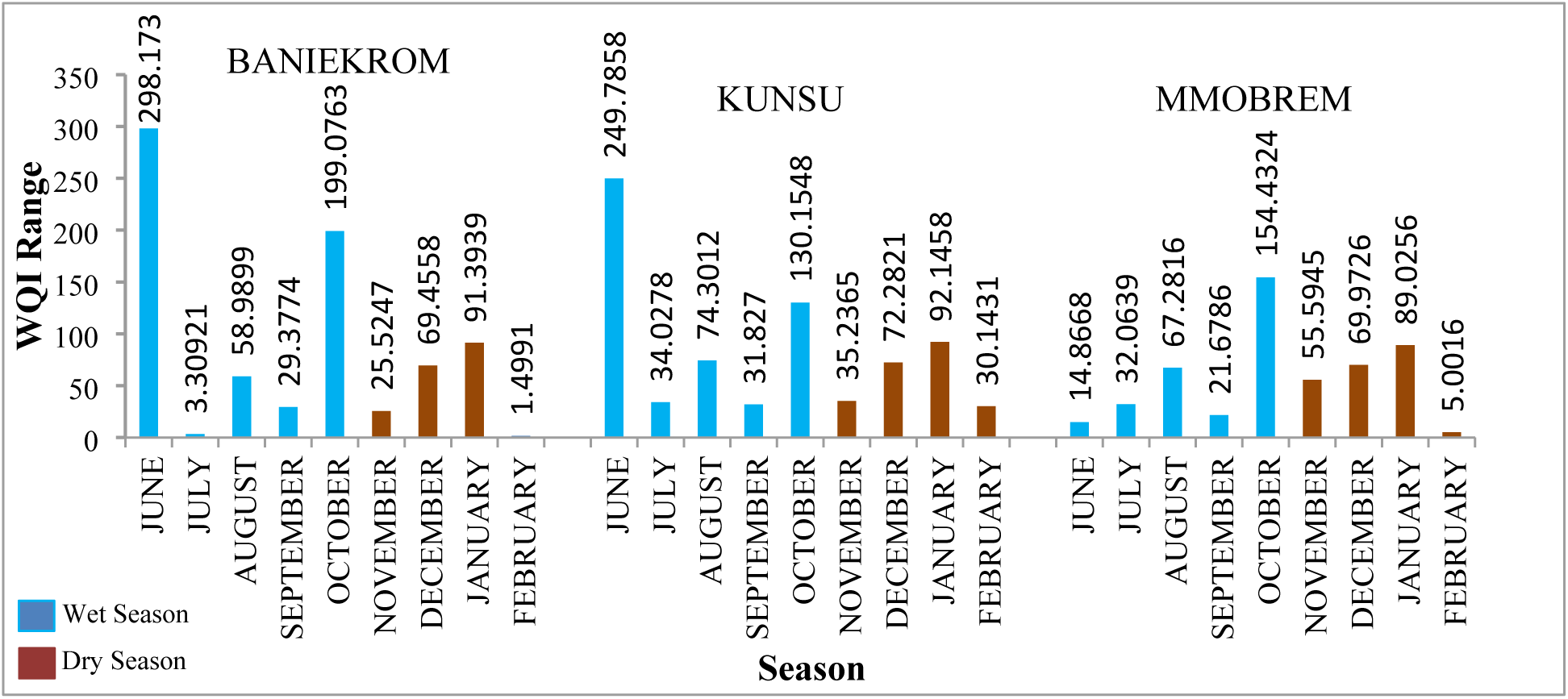
Water Quality Index for Baniekrom, Kunsu & Mmobrem

The mean WQI for Kunsu was 97.49 in the wet season and 71.99 in the dry season, indicating lower water quality during the dry period. Monthly WQI values ranged from 249.79 to 31.83 (June–September) in the wet season and 130.15 to 30.14 (October–February) in the dry season (Fig. 4), showing notable temporal variation. These results suggest that while wet-season rainfall improves water quality through dilution, some dry-season periods may render the water unsuitable for drinking without treatment, emphasizing the need for ongoing monitoring and management.

For Mmobrem, the mean WQI was 33.97 in the wet season and 74.81 in the dry season, indicating lower water quality during the wet season and improvement in the dry season. Monthly WQI values ranged from 5.00 to 154.43 between June and February, reflecting substantial temporal variability. The low wet-season values suggest that surface water may be unfit for drinking without treatment, while the higher dry-season WQI indicates improved quality, likely due to reduced runoff and pollutant inputs.

### 3.6 Hydrochemical Facies as an Integrative Indicator of Land-Use Impact on Surface Water Quality

Hydrochemical characterization of surface waters using Piper trilinear diagram analysis

Hydrochemical characterization of surface waters within the micro-watershed, using Piper trilinear diagram analysis, is presented in Figs. 5-7. Fig. 5 is a Piper plot for the Mmobrem study area, Fig. 6 is for Kunsu, and Fig. 7 is for Baniekrom. Piper plot diagram for Mmrobem water samples illustrated the hydrochemical facies at the Mmrobem site (Fig. 5). Sample points cluster in the Calcium-Chloride (Ca-Cl) field of the central diamond, with a strong anion dominance toward chloride (Cl⁻) in the lower triangle. For Kunsu, the piper plot diagram pointed to a clear and strong clustering in the Calcium-Chloride (Ca-Cl) type field, with a pronounced shift toward the chloride (Cl⁻) apex in the anion triangle (Fig. 6). The Piper plot diagram for the Baniekrom site indicates a mixed Ca-Mg-Na+K cation field, with a clear dominance of Chloride (Cl⁻) in the anion triangle. This results in a classification of Ca-Cl to mixed-type in the central diamond (Fig. 7).

**Fig. 5.**
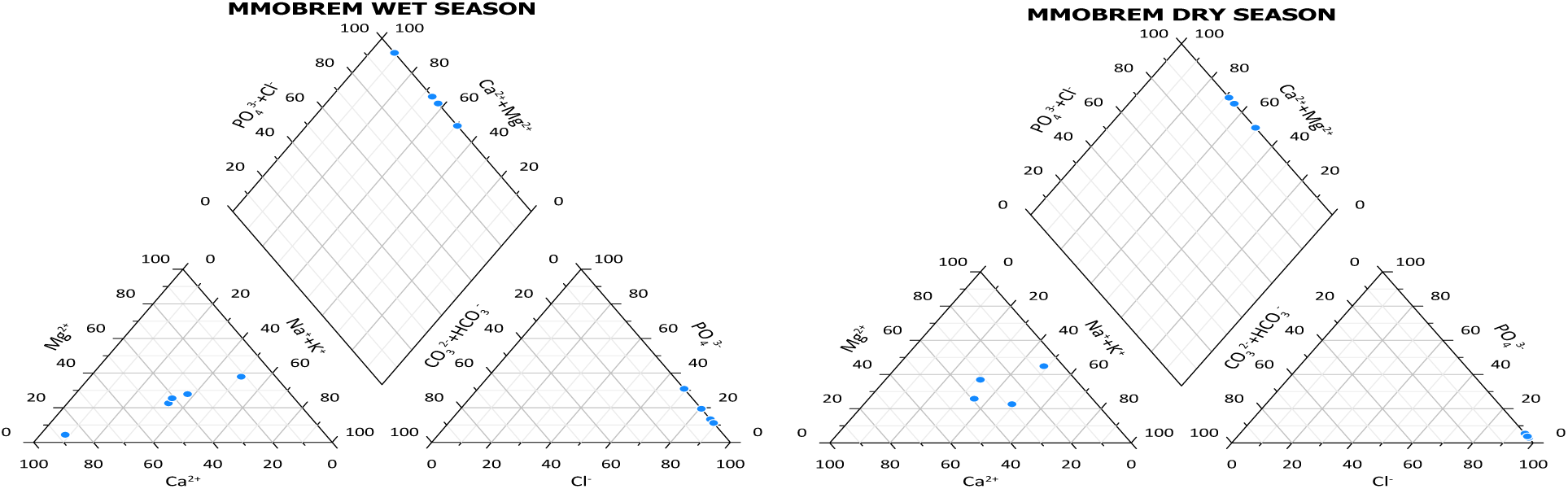
Generate Piper Plot Diagram for MMBREM Water Samples

**Fig. 6.**
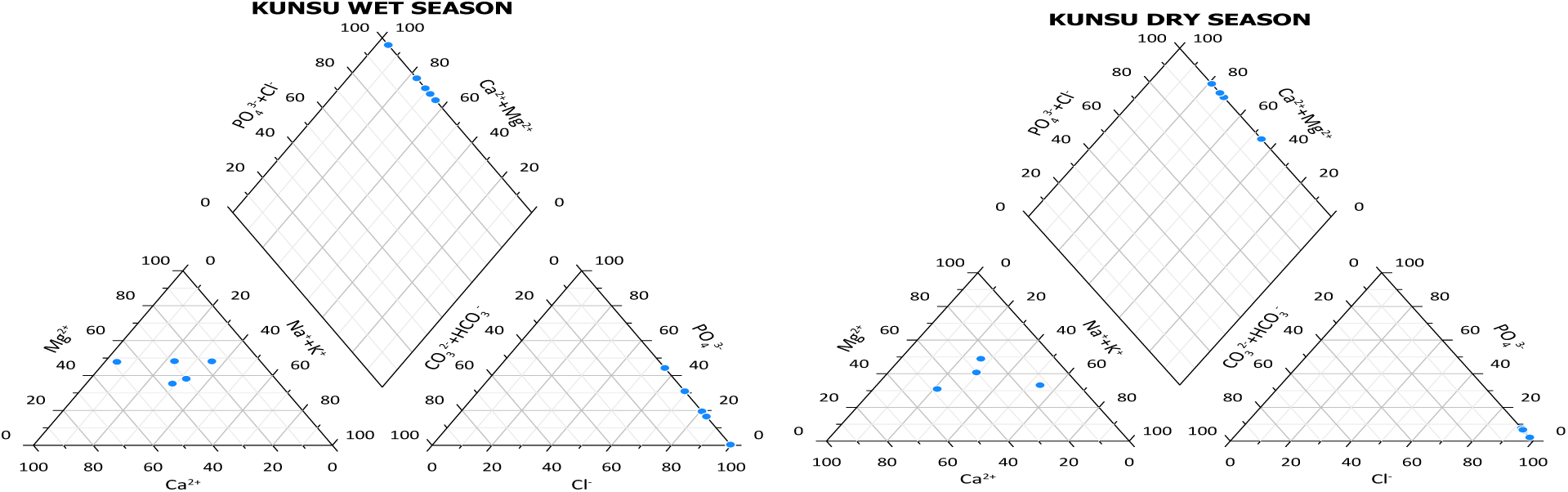
Generated Piper Plot Diagram for KUNSU Water Samples

**Fig. 7.**
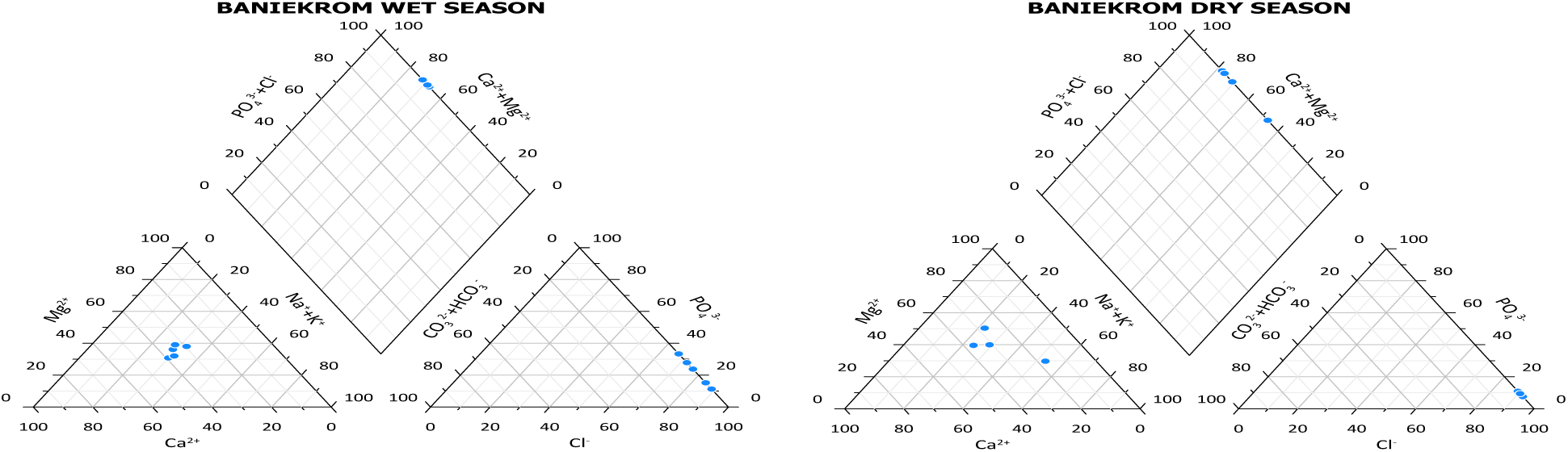
Generated Piper Plot Diagram for BANIEKROM Water Samples

### 3.7 Ecological Risk Assessment

In aquatic ecosystems, the ecological risks posed by various elements differ depending on the species and the specific element in question. Calcium, magnesium, sodium, and potassium exhibit a higher ecological risk when it comes to their impact on algae (Table 3). This elevated risk is due to their potential to disrupt the delicate balance of nutrients and ions in aquatic environments, which can lead to negative consequences for algae populations. On the other hand, elements such as zinc, cadmium, and copper present a relatively lower risk to algae. Although potentially harmful in large quantities, these elements do not appear to exert the same level of ecological pressure as calcium, magnesium, sodium, and potassium in typical environmental concentrations. When assessing the impact of these elements on daphnia, a type of small aquatic crustacean, the overall risk is notably lower than that observed in algae. Daphnia, known for their sensitivity to changes in water quality, seem to be less affected by the presence of these elements, indicating a minimal ecological risk. In fish populations, the risk posed by calcium, magnesium, sodium, and potassium remains minimal. Since fish are higher up in the aquatic food chain, they are less directly affected by the concentrations of these elements, which are more likely to impact primary producers, such as algae. Consequently, the ecological risks associated with these elements for fish are significantly lower, indicating that they do not pose a substantial threat to fish populations in most aquatic ecosystems.

**Table 3.** Ecological risk output.

| **Metal** | **MEC**<br>**(mg/L)** | **PNEC**<br>**algae**<br>**(mg/L)** | **RQ**<br>**algae** | **PNEC**<br>**daphnid**<br>**(mg/L)** | **RQ**<br>**daphnid** | **PNEC**<br>**fishes**<br>**(mg/L)** | **RQ**<br>**fishes** |
| --- | --- | --- | --- | --- | --- | --- | --- |
| **Zn** | 0.1065 | 1202.333 | 8.86E-05 | 4145.877 | 2.57E-05 | 9172.84 | 1.16E-05 |
| **Cd** | 0.0024 | 1059.876 | 2.26E-06 | 3137.088 | 7.65E-07 | 6689.198 | 3.59E-07 |
| **Fe** | 11.9512 | 1606.832 | 7.44E-03 | 6213.015 | 1.92E-03 | 14132.52 | 8.46E-04 |
| **Cu** | 0.8448 | 1329.335 | 6.36E-04 | 4762.184 | 1.77E-04 | 10634.17 | 7.94E-05 |
| **Ni** | 0.1139 | 1227.822 | 9.28E-05 | 4398.527 | 2.59E-05 | 9822.102 | 1.16E-05 |
| **Co** | 0.0375 | 344.4659 | 1.09E-04 | 909.239 | 4.12E-05 | 1885.798 | 1.99E-05 |
| **Pb** | 0.4606 | 545.8534 | 8.44E-04 | 1190.441 | 3.87E-04 | 2357.626 | 1.95E-04 |
| **Mn** | 1.3367 | 321.1141 | 4.16E-03 | 847.6006 | 1.58E-03 | 1757.958 | 7.60E-04 |
| **Cr** | 0.1922 | 303.9184 | 6.32E-04 | 802.2114 | 2.40E-04 | 1663.819 | 1.16E-04 |
| **Na** | 20.0166 | 661.4872 | 3.03E-02 | 2557.721 | 7.83E-03 | 5817.956 | 3.44E-03 |
| **K** | 21.4375 | 1124.97 | 1.91E-02 | 4349.833 | 4.93E-03 | 9894.406 | 2.17E-03 |
| **Mg** | 14.053 | 550.6157 | 2.55E-02 | 1972.515 | 7.12E-03 | 4404.712 | 3.19E-03 |
| **Ca** | 19.7101 | 880.5751 | 2.24E-02 | 3154.555 | 6.25E-03 | 7044.26 | 2.80E-03 |
| **As** | 0.0011 | 222.3759 | 4.98E-03 | 494.3216 | 2.24E-03 | 983.5166 | 1.13E-03 |
| **Hg** | 0.0061 | 628.7026 | 9.70E-04 | 1429.385 | 4.26E-04 | 2859.482 | 2.13E-04 |

## 4. DISCUSSION

The findings of this study underscore the significant impact of anthropogenic activities on the surface water quality of the Ahafo Ano Southwest District. The elevated levels of various pollutants, including heavy metals, nitrate, phosphate, and total suspended solids, are clear indicators of the detrimental effects of human activities on the environment. These findings are consistent with previous studies conducted in similar regions, highlighting the pervasive nature of water pollution in developing countries [4,6,12]

The correlation analysis revealed significant relationships between different pollutants, suggesting that the presence of one pollutant may influence the concentration of others. For instance, the strong positive correlation between total suspended solids and copper indicates that increased levels of suspended solids may contribute to the accumulation of copper in the water bodies. This finding is supported by previous research, which has demonstrated the role of suspended sediments in the transport and deposition of heavy metals [23]. Principal component analysis identified distinct clusters of observations, indicating that variations in water quality are influenced by multiple factors, including anthropogenic activities such as mining, agriculture, land use patterns, and population density. Further research is needed to elucidate the specific drivers of water quality changes in the study area. Analysis of seasonal variation shows elevated Zn, Cd, and Fe during the wet season, likely linked to runoff from mining activities, while higher Cu concentrations in the dry season may reflect agricultural inputs or localized domestic sources. The spatial distribution of contaminants in the Mankran watershed reflects dominant land-use activities in the Ahafo Ano Southwest District. Elevated Cu and Pb are linked to artisanal and small-scale gold mining, where ore processing and tailings disposal introduce metal-rich sediments into streams, especially during the wet season. In contrast, nitrate and phosphate enrichment is associated with agricultural activities such as fertilizer use, livestock rearing, and runoff from farmlands, with higher wet-season levels driven by increased surface runoff. The Ca–Cl hydrochemical facies indicates geogenic control from rock weathering, with additional chloride inputs likely from agricultural and domestic sources, reflecting combined natural and anthropogenic influences [24]. pH levels were slightly higher in the dry season, indicating more alkaline conditions, while increases in TDS and EC likely reflect evaporation and concentration of dissolved ions from mining and agricultural runoff. Elevated TSS during the dry season suggests increased particulates, potentially from soil erosion associated with land use and mining activities [24]. This observation may be linked to anthropogenic activities, such as mining, as decreased precipitation during the dry season could lead to higher TSS levels. Nitrate levels showed a substantial increase in the wet season, while Phosphate levels exhibited a slight rise during the same period compared to the dry season. These findings point to a heightened influx of certain metals and nutrients during the wet season, ascribed to intensified agricultural use, while the dry season potentially leads to more outstanding suspended particulate matter in the water, possibly due to mining activities.

The ecological risk assessment revealed that certain elements, such as calcium, magnesium, sodium, and potassium, pose higher risks to algae, while others, like zinc, cadmium, and copper, have lower ecological risks which could be due to differences in species sensitivity. These findings are consistent with previous studies that have investigated the toxicity of different elements to aquatic organisms [25].

The study’s findings have important implications for water resource management in the Ahafo Ano Southwest District. The elevated levels of pollutants pose significant risks to aquatic ecosystems and human health. Effective measures are needed to mitigate the adverse effects of human activities and promote sustainable water use. This may include implementing stricter regulations for industrial and agricultural activities, promoting sustainable land use practices, and investing in wastewater treatment infrastructure.

The seasonal analysis of the Water Quality Index (WQI) across Baniekrom, Kunsu, and Mmobrem revealed distinct spatial patterns that are strongly influenced by land use activities, specifically agriculture, deforestation, and mining within each catchment. The results demonstrate pronounced differences between the wet and dry seasons, reflecting the interaction between hydrological conditions and pollutant transport processes. Baniekrom recorded mean WQI values of 97.462 in the wet season and 77.39 in the dry season, indicating that water quality generally deteriorates during rainfall periods. The wet season WQI values, particularly the peak of 298.17, suggest episodes of severe contamination. This deterioration is most likely driven by intense surface runoff, which transports fertilizers, pesticides, animal waste, and eroded sediments from farmlands and deforested areas directly into the river [26]. Deforestation typically exposes soil surfaces, enhancing erosion rates during storms, which subsequently elevates turbidity and nutrient loads in surface waters. In contrast, the lower dry-season values (minimum 1.50) reflect the absence of runoff-driven inputs, resulting in relatively improved water conditions. Kunsu exhibited mean WQI values of 97.485 (wet season) and 71.992 (dry season), showing a similar seasonal trend as Baniekrom, with poorer water quality during the wet months. The highest wet season WQI value (249.79) indicates substantial degradation likely linked to small-scale mining activities within the area. During the wet season, runoff can easily mobilize mine tailings, heavy metals, suspended particulates, and acidic drainage products from exposed mining surfaces into the stream. These processes intensify during rainfall, resulting in elevated levels of contamination. The dry season WQI values (down to 30.14) suggest reduced pollutant transport, as rainfall-driven flushing diminishes and fewer sediments or metals are washed into the water system. Unlike Baniekrom and Kunsu, Mmobrem displayed an opposite seasonal pattern. The wet-season mean WQI (33.971) was significantly lower than the dry-season mean (74.805), indicating comparatively better water quality during the rainy period. Wet-season WQI values remained low (e.g., 14.87, 21.68) except for an isolated maximum of 154.43, suggesting that although runoff introduces pollutants, the overall effect is overshadowed by strong dilution from high rainfall volumes [27]. This dilution reduces the concentration of dissolved solids, nutrients, and other contaminants, leading to improved wet season water quality. Conversely, during the dry season, reduced streamflow and increased evaporation promote the concentration of pollutants. Continued agricultural input without adequate flushing may also contribute to the elevated dry season WQI values. The observed improvement in Mmobrem’s wet-season WQI is attributed to rainfall-driven dilution. While direct gauging or discharge data were not available for this study, this interpretation is supported by regional hydrological patterns and precipitation records indicating increased flow during the wet season, which would dilute pollutants and reduce contaminant concentrations. The Piper diagram highlights the dominant hydrochemical facies and underlying geochemical processes controlling groundwater chemistry. While natural hydrogeochemical reactions shape the overall composition, chloride enrichment in several samples suggests additional anthropogenic influences, with elevated levels likely arising from agricultural runoff (fertilizers and irrigation return flows) and domestic sources (sewage or wastewater). The distinct facies classification therefore reflects a combination of natural mineral dissolution and external human inputs, providing a comprehensive view of factors affecting groundwater quality in the study area.

The cation plot shows most samples fall within the Ca–Mg–Na+K dominance field, indicating control by alkaline earth and alkali metals. While Basalt weathering contributes naturally to these minerals, elevated Na⁺ and K⁺ levels may also reflect anthropogenic inputs from agricultural runoff and mining activities, suggesting both geogenic and human influences on mineral enrichment [28].

In the anion triangle, the samples cluster predominantly in the chloride (Cl⁻) dominant zone, indicating that Cl⁻ is the principal anion in the system. Chloride-rich waters are often linked to prolonged water–rock interactions, evaporation effects, or external inputs such as anthropogenic contamination, atmospheric deposition, or intrusion from deeper saline formations [29]. The strong Cl⁻ dominance observed may therefore reflect the influence of evaporative concentration, domestic/agricultural inputs, or geogenic chloride sources, depending on local geology and land use activities.

When projected into the central diamond field, the samples plot within the Ca–Cl facies, corresponding to an alkaline earth–chloride water type. These facies are typically associated with relatively evolved groundwater that has undergone significant mineral dissolution along its flow path [30].Water of this type often indicates advanced geochemical evolution, which may involve ion exchange, dissolution of evaporite minerals, or mixing with saline or brackish water [31]. The position of the samples in the upper right sector of the diamond further suggests increasing total dissolved solids (TDS) and higher mineralization, characteristics commonly found in water from semi-arid regions or areas with long residence time [32].

Overall, the Piper diagram results indicate that groundwater in the study area is characterized by a Ca–Cl hydrochemical facies, reflecting a combination of carbonate dissolution, chloride enrichment, and possible anthropogenic or natural salinization processes. This facies distribution highlights the importance of both geogenic controls and external inputs on the surface water chemistry. The dominance of alkaline earths and chloride suggests that the water system may be vulnerable to mineralized recharge, evaporative concentration, or human activities that contribute chloride to the system, such as agriculture, waste disposal, and urban runoff. This study is limited by the absence of streamflow data, restricting the assessment of dilution effects, particularly at Mmobrem. Seasonal sampling may not capture short-term variability, and the lack of direct source measurements limits precise attribution of pollutants. The findings emphasize the need for integrated watershed management aligned with the Environmental Protection Agency Ghana and the Ghana Minerals Commission. Key measures include stricter regulation of mining activities, adoption of agricultural best management practices, and implementation of riparian buffer zones, supported by improved monitoring and community education.

## 5. CONCLUSION AND RECOMMENDATION

This study presents a comprehensive assessment of the surface water quality in the Ahafo Ano Southwest District, highlighting significant pollution levels due to anthropogenic activities. Elevated concentrations of heavy metals, nitrate, phosphate, and total suspended solids pose substantial risks to aquatic ecosystems and human health. The correlation analysis reveals interconnectedness among pollutants, emphasizing the need for integrated management approaches. Seasonal variations in pollutant levels, particularly the increased influx of nutrients during the wet season and elevated suspended solids during the dry season, underscore the complex interplay between natural factors and human activities. The ecological risk assessment identifies certain elements as having higher risks to algae, while others pose lower risks to daphnia and fish.

The seasonal assessment of water quality across Baniekrom, Kunsu, and Mmobrem reveals that hydrological variability and land-use activities exert significant control over the chemical and ecological condition of the streams. The pronounced wet-season deterioration observed in Baniekrom and Kunsu underscores the strong influence of surface runoff, which mobilizes agricultural residues, eroded sediments, and mining-derived contaminants into the water bodies. In contrast, the improved wet-season water quality in Mmobrem highlights the role of dilution during high flows, despite localized contamination events. These findings emphasize the sensitivity of catchments with intensive agriculture, deforestation, and mining to rainfall-driven pollutant transport.

Hydrochemical facies derived from the Piper diagram further support the interaction between geogenic and anthropogenic factors in shaping water chemistry. The dominance of Ca–Cl type water suggests significant contributions from basaltic weathering, chloride enrichment, and mineral dissolution processes, in addition to external inputs from agriculture and domestic sources. The clustering of samples within high TDS zones indicates increasing mineralization and the progressive evolution of water chemistry along flow paths. Overall, the integration of WQI and hydrochemical facies analysis demonstrates that the study area is characterized by both natural geological controls and substantial human-induced impacts, with water quality varying markedly between seasons. These results highlight the need for continuous monitoring, sustainable land-use management, and mitigation strategies particularly during the wet season to safeguard water resources for domestic, agricultural, and ecological use. The findings underscore the pressing need for sustainable water management practices to safeguard the environment and ensure the well-being of future generations. Further research is necessary to deepen our understanding of the factors driving changes in water quality and to develop practical solutions for addressing these challenges.

## Availability of data and materials

All raw and processed data generated or analyzed during this study are included in this published article.

## Ethics approval and consent to participate

Not applicable; no human or animal subjects were directly used in the study for which approval ought to be sought.

## Consent for publication

All authors have proofread the manuscript and approved the submission.

## Credit authorship contribution statement

David Azanu was in charge of drafting, Afia Sarpong Anane Gyebi collected and analyzed the samples, Simeon Makafui Nutsuklo, Sampson Yaw Marfo and Isaac Ayew Aidoo assisted in analysing the samples, Francis Nyarkoh and Evans Barimah assisted in data analysis and writing, Seifu A Tilahun and Junias Adusei-Gyamfi were in charge of research conceptualization, Methodology, oversight, and leadership.

## Declaration of Competing Interest

The authors declare that they have no known competing financial interests or personal relationships that could have appeared to influence the work reported in this paper.

## Acknowledgements

We are grateful to the SHEATHE Project (www.sheathe.org) hosted by the Department of Chemistry, KNUST, Kumasi, Ghana, for using their laboratory resources and Royal Roads University, Canada, for instrumentation.

## Notes

### Competing Interest Statement

The authors have declared no competing interest.

